# Emu: Species-Level Microbial Community Profiling for Full-Length Nanopore 16S Reads

**DOI:** 10.1101/2021.05.02.442339

**Authors:** Kristen D. Curry, Qi Wang, Michael G. Nute, Alona Tyshaieva, Elizabeth Reeves, Sirena Soriano, Enid Graeber, Patrick Finzer, Werner Mendling, Qinglong Wu, Tor Savidge, Sonia Villapol, Alexander Dilthey, Todd J. Treangen

## Abstract

16S rRNA based analysis is the established standard for elucidating microbial community composition. While short read 16S analyses are largely confined to genus-level resolution at best since only a portion of the gene is sequenced, full-length 16S sequences have the potential to provide species-level accuracy. However, existing taxonomic identification algorithms are not optimized for the increased read length and error rate of long-read data. Here we present Emu, a novel approach that employs an expectation-maximization (EM) algorithm to generate taxonomic abundance profiles from full-length 16S rRNA reads. Results produced from one simulated data set and two mock communities prove Emu capable of accurate microbial community profiling while obtaining fewer false positives and false negatives than alternative methods. Additionally, we illustrate a real-world application of our new software by comparing clinical sample composition estimates generated by an established whole-genome shotgun sequencing workflow to those returned by full-length 16S sequences processed with Emu.

## Introduction

Sequencing the 16S subunit of the ribosomal RNA gene has been a reliable way to characterize diversity in a community of microbes since Carl Woese used this technique to identify Archaea in 1977^1^. Today, high-throughput sequencing machines used for this analysis are dominantly Illumina devices. Although cost-effective and accurate, Illumina sequencers are limited to joined paired-end reads of roughly 500 nucleotides (bps). Since the 16S gene is approximately 1,550 bps, analysis is then restricted to target amplification of just a subset of the hypervariable regions. This prevents distinction between highly similar species, and ultimately produces taxonomic profiles that can in most cases only be measured down to the genus level.

Recent developments in sequencing technology, from providers like Pacific Biosciences and Oxford Nanopore Technologies (ONT), permit amplification of sequences spanning the entire 16S gene. However, these long-reads come with one notable drawback: high rates of sequencing error. While Illumina reads contain no more than one error per 100 base pairs, long-read sequences yield errors in 10-15% of the nucleotide positions.

The canonical pipeline for 16S analysis operates in two main steps. First, the set of raw sequences is de-noised to identify a smaller set of core sequences, where each set is believed to represent a distinct taxonomic unit in the community. Various algorithms are available for this process^2^, yet all are calibrated to the level of error associated with Illumina reads. Second, the representative sequences are compared to a database and assigned a taxonomic label. Since reads are already corrected for error at this point, a database lookup tool such as BLAST^3^ is effective here. Yet, the error-correction presented by these pipelines are simply not designed for error-rates of 10-15% produced by current long-read sequencers. Thus, these pipelines are not able to produce accurate results from such reads.

Since ONT sequencers are comparatively recent to the marketplace, 16S method development for these devices has only just begun^4^. In the absence of dedicated tools, some publications have chosen to use a more general read-mapping software such as BWA-MEM^5^ or the LAST aligner^6^ to align reads directly to raw 16S sequences in one of the major databases^7–9^. Another approach involves use of Kraken 2^10^ and its Bayesian cousin Bracken^11^, since this combination compared favorably to the default classifier in QIIME 2^12^ for 16S short reads.

NanoClust^13^ became the first published method purpose-built for taxonomic abundance profiling using full-length 16S amplicon sequencing from ONT machines. As its name implies, this method follows steps that resemble the de-noising/database-lookup procedure discussed above. Here, the two-stage procedure is implemented in Nextflow^14^ and uses external tools for demultiplexing, quality-filtering, clustering, polishing and taxon assignment. As with Illumina software, taxon assignment is lightweight since this step includes only a small number of ostensibly high-quality representative sequences. Yet due to use of consensus sequences, this method is susceptible to overlook identification of species that are truly present in the error-prone dataset.

In terms of classification from shotgun long sequences, Centrifuge^15^ and MetaMaps^16^ are commonly used. Centrifuge, although not designed specifically for long or high-error reads, was tested on a set of ONT shotgun sequencing reads in its publication. The results show it to be an effective tool for this purpose, with particular emphasis on recall when a minimum threshold of 5 mapped reads is applied to the species identified. ONT now includes Centrifuge as a step in WIMP^17,18^ (What’s in my Pot?), a long-read 16S workflow provided on its EPI2ME analysis platform.

MetaMaps *was* designed for long, high-error reads and is intended for taxonomic binning of shotgun sequencing reads. In order to address the error profile in the data, MetaMaps uses an approximate read-mapping algorithm to identify multiple candidate species and locations for each read. It then applies an expectation-maximization (EM) algorithm to adjust the relative confidence in each mapping based on the mapping density of other reads in the sample. This has the effect of smoothing out some of the noise that is inherently created by the ONT error profile, resulting in a more accurate relative abundance estimation. While effective for whole-genome sequences, the approximate mapping algorithm in MetaMaps does not achieve the required resolution for 16S analysis and is thus not suitable for this scenario. However, this and other EM algorithms that have been used previously in related settings for disambiguating ambiguous read mappings^19, 20^, provoke interest in an EM method for error-correction of long 16S reads.

Here, we present Emu, a microbial community profiling software tool tailored for full-length 16S data with high error rates. It capitalizes on the benefits of increased read length, while incorporating a crucial error-correction step. Emu’s algorithm involves a two-stage process. First, proper alignments are generated between reads and the supplied reference database. Emu then implements an EM-algorithm to iteratively refine species-level relative abundances based on total read-mapping counts. This results in microbial community profile estimations from full-length 16S reads which are more accurate than existing methods at both the genus and species level.

## Results

To demonstrate the performance of Emu, two studies were completed. First, a quantitative comparison using three distinct communities, where an actual or *de facto* ground truth could be used for evaluating accuracy and comparing methods. Second, a demonstration of Emu’s applicability to understanding dynamics in actual microbial communities. In this real-world model, human vaginal microbiome clinical samples were processed with two separate pipelines: 16S long reads analyzed with Emu, and whole genome shotgun sequences processed with Bracken.

### Quantitative comparison

To quantify the output of Emu in relation to several existing methods, three communities were used. The first is a single data set of simulated ONT reads which follows the distribution of a published mock community. The other two are synthetic mock communities, each of which were sequenced with both ONT and Illumina devices. Performance of each method was evaluated at both the genus and species level using three metrics: the L1-norm of the taxonomic abundance profile, the count of true positive taxa (TP), and the count of false positive taxa (FP). Computational resources required by each method were measured by recording the run time and memory usage for each software.

The set of methods used for comparison include those discussed above: Kraken 2, Bracken, NanoClust, and Centrifuge. We also include QIIME 2^12^ and the primary alignment generated by minimap2^21^. Although minimap2 is not a composition estimator or read-level classifier in itself, it is included because it is instrumental in the Emu algorithm: mnimap2 is the read-mapping software Emu uses to compute likelihood scores and iteratively estimate relative abundance. Including minimap2 in the comparison separates the effect of the EM impelementation in Emu from the read-mapping output it uses as a starting point. QIIME 2 is a suite of software with high adoption primarily used with Illumina sequencing data, and is included mainly to illustrate the efficacy of ONT reads with a method trusted on Illumina reads. Identical reference databases were built for each software to ensure even comparison across methods (see **Methods** for details).

Ground truth relative abundance values for synthetic communities are based on sample-specific imputed values rather than on estimated CFU-counts during production. This was done to correct for fluctuations in true abundance which may occur during handling, storage or library preparation (including potential primer bias during 16S amplification). The two ZymoBIOMICS community profiles are reasonably similar to their abundance claims, but the synthetic gut community is subject to greater variation by nature of the microbes included and the skewed distribution. Therefore, this dataset-specific imputed ground truth is necessary to reliably evaluate community composition. Details on this process are described in the Establishing ground truth section in **Methods** below, and for these two communities the term ground truth herein refers to this imputed value.

#### MBARC-26 simulated mock community dataset

ONT reads were simulated following the composition of a published mock community, MBARC-26^22^, which contains 23 bacterial and 3 archaeal strains. Detailed information on the reference sequences and distribution of simulated reads are contained in Supplementary Table 1.

#### ZymoBIOMICS mock community standard dataset

A previous study compared 16S sample composition accuracy across a series of hypervariable regions as well as the full-length gene using the ZymoBIOMICS community standard catalog number D3605^23^. We retrieved the ONT full-length dataset and one of the Illumina datasets for our analysis. We selected the Illumina dataset with targeted regions V4-V6 to represent short-read data since the study showed this dataset to produce classification results among the most accurate for this community specifically.

#### Synthetic mock gut microbiome dataset

Finally, to challenge our software, a synthetic community mimicking the human gut microbiome was created and sequenced with both ONT and Illumina devices, as described in the Gut Microbiome Mock Community Sample Creation section in **Methods**. To represent a real-world scenario with unknown species, *Romboutsia hominis* is included in the sample, even though this new species is not present in our database. The derived relative abundances of 21 species present in the sample are described in Supplementary Table 1. One notable difference between the two datasets for this community is that *Bifidobacterium dentium* is not considered to be a True Positive in the ONT sequences. This is a result of a recently noted issue with the standard ONT forward primer, which contains three mismatching bases to the family Bifidobacteriaceae and thus fails to amplify microbes of this taxa^8^. As a result the ONT data do not contain reads from this microbe, so considering it a true positive for this data set is inaccurate. This is one example of why an imputed ground truth was used for the mock communities. This situation also highlights the need for additional research into primers in this region to identify reliably universal primers or characterize any sensitivity gaps.

### Performance

Results of all methods on the simulated data set and two synthetic mock communities are contained in Table 1. Computational resources required by each method are listed in Supplementary Table 2. Complete abundance profile output from all methods on all data sets are provided in Supplementary File 2. All results generated utilize the default Emu database.

**Table 1.**
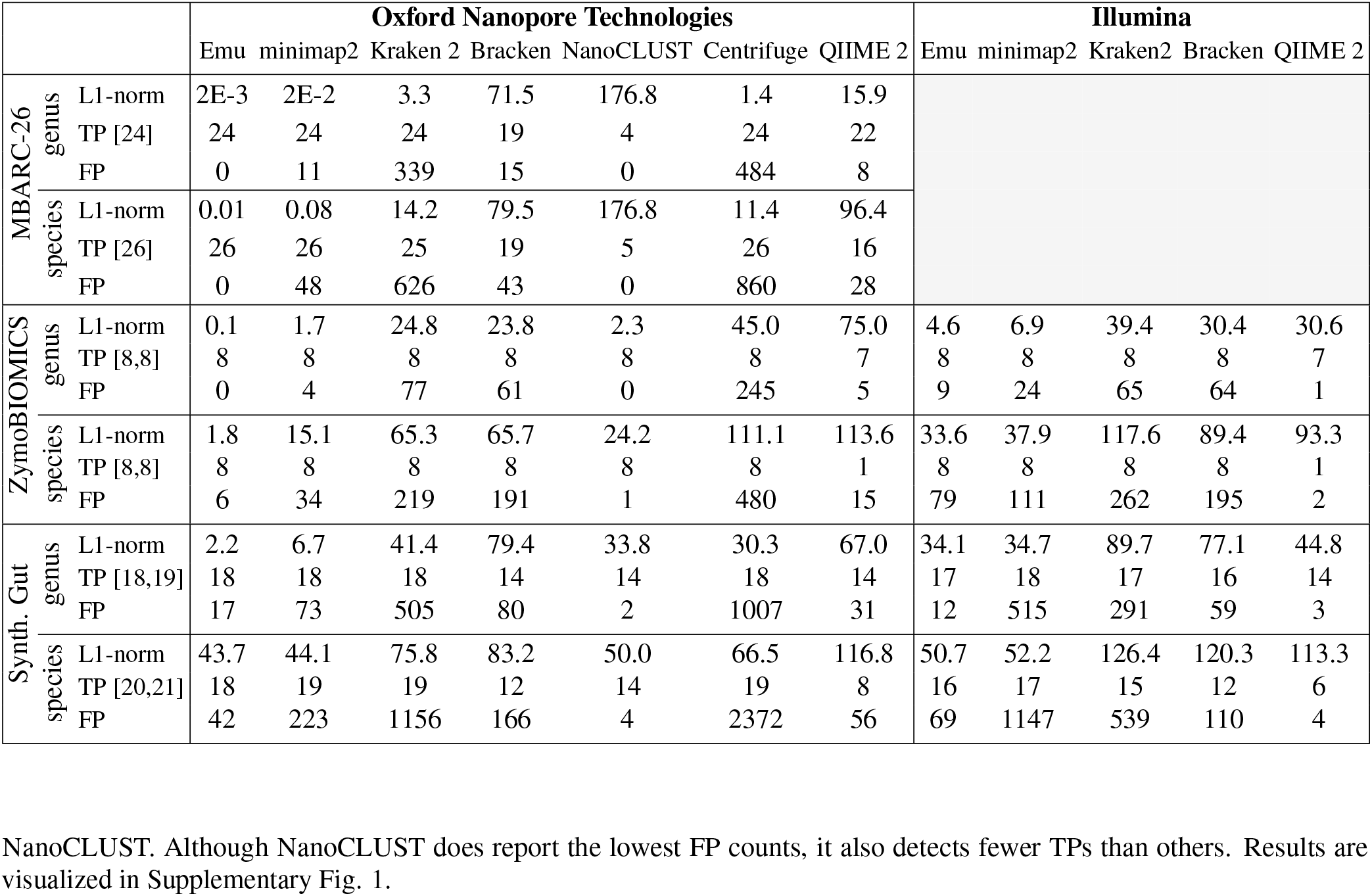
Performance summary of 16S relative abundance estimates on ONT and Illumina sequences for all three communities. The row headers for TP contain the estimated number of actual true positives for each data set: for [*x, y*]: *x* denotes the expected TP for Nanopore data set, and *y* for Illumina data set.

#### MBARC-26 simulation

For the MBARC simulated data, we see in Fig. 1 that Emu outperforms every method. Not only does Emu express the lowest L1 distance, but it is also the only method to correctly identify all 26 species without producing any false positives. Notably, there is a substantial difference in both L1 distance and false positive counts between Emu and minimap2. This reflects the accuracy gains produced by the EM algorithm compared to a simple similarity-based taxon assignment approach. It is evident from the memory and run time data between these two methods (Supplementary Table 2) that the majority of computation resources used by Emu are in fact due to its use of minimap2 for alignment generation. NanoCLUST results differ from other methods shown in that it has no false positives, but fails to identify several of the present taxa; in other words, it is generally conservative in its identifications.

**Figure 1.**
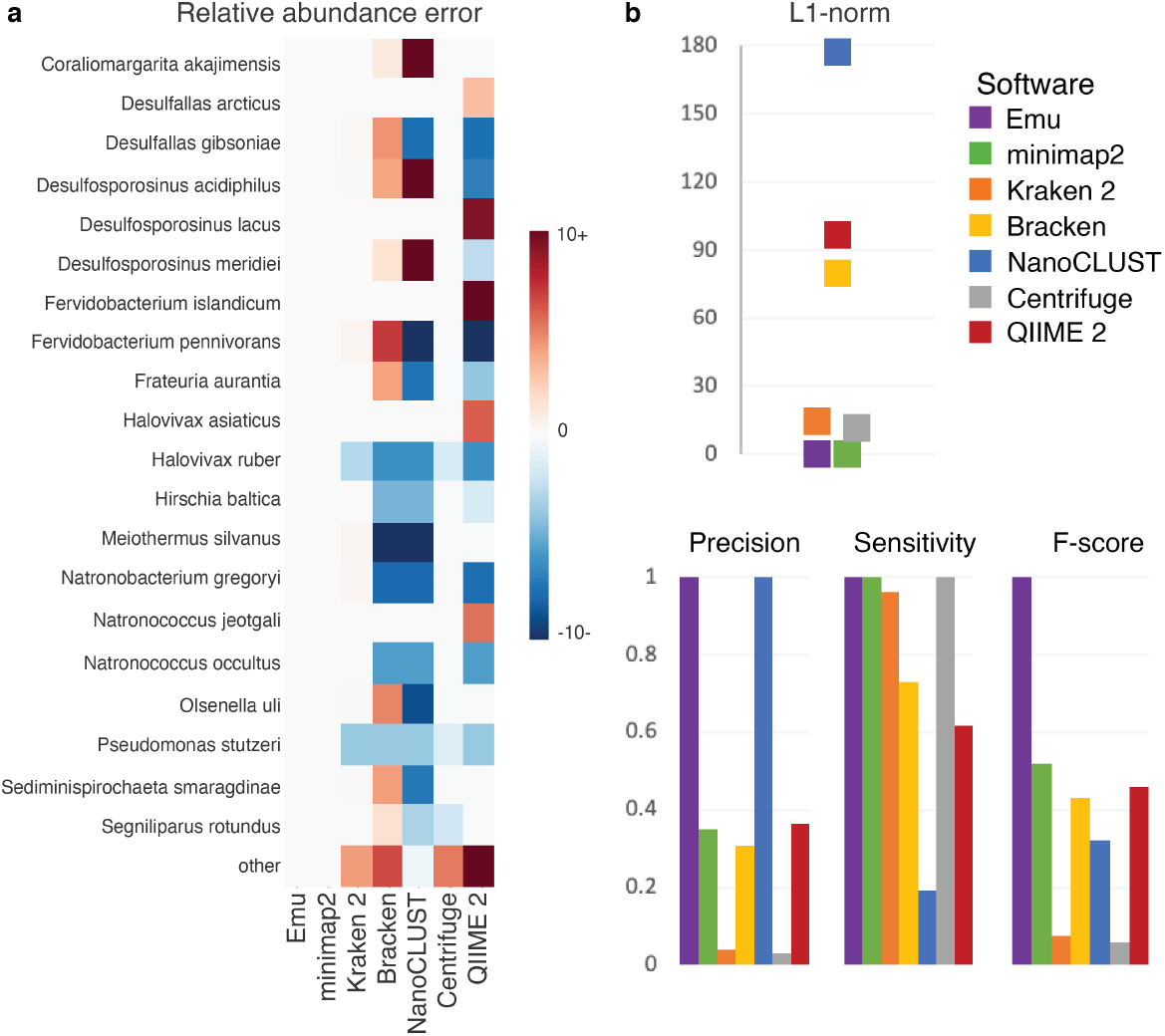
Performance on simulated nanopore reads (MBARC). **(a)** Species-level error between theoretical and estimated relative abundances, where darker blue denotes an underestimate by the software, darker red denotes an overestimate, and white represents no error. Color scheme is capped at ±10, resulting in error greater than ±10% observing the maximum error colors. Displayed are the 20 species claiming the largest abundance in any of the Emu, minimap2, Kraken 2, Bracken, NanoCLUST, Centrifuge, or QIIME 2 results. “Other” represents the sum of all species not shown in figure for the respective column. **(b)** Species-level L1-norm, precision, sensitivity, and F-score for the seven methods evaluated in panel **a**.

#### ZymoBIOMICS

Emu on ONT reads expresses the lowest L1 distance across the methods tested at both the genus and species levels. While almost all methods accurately detect the 8 species in the sample, the number of false positives reported varies. Of the methods with perfect sensitivity, NanoCLUST returns the fewest false positives and Emu returns the second fewest. It is also important to note that the abundance accuracy and sensitivity measured in the ONT dataset proves superior to those of the Illumina dataset, especially at the species level. When restricting to only the Illumina results, Emu again proves the lowest L1 distance. While we do not recommend using Emu on Illumina 16S reads, this shows Emu to be a sensible approach regardless of the read-error profile. Fig. 2 provides a graphical representation of accuracy measures.

**Figure 2.**
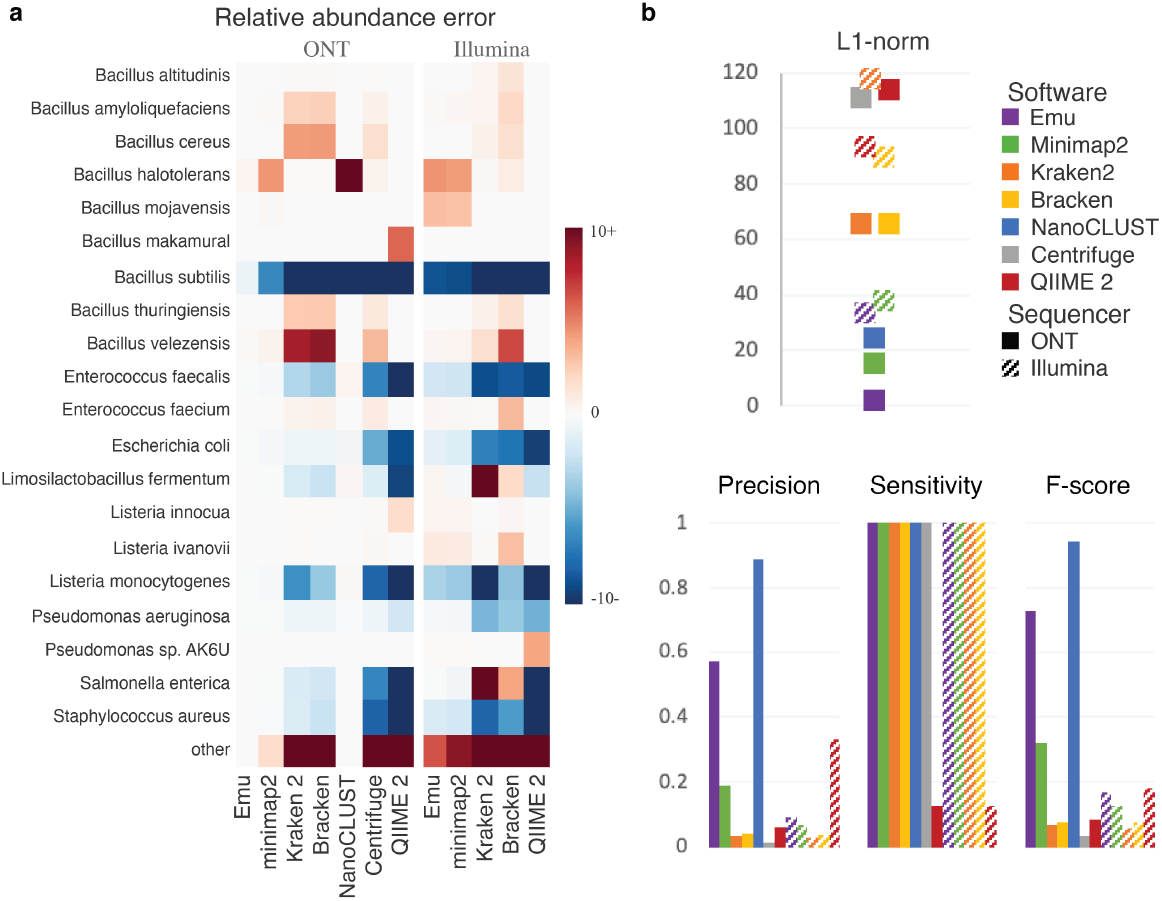
Performance on ZymoBIOMICS community standard. **a** Species-level error between calculated ground truth and estimated relative abundances, where darker blue denotes an underestimate by the software, darker red denotes an overestimate, and white represents no error. All Oxford Nanopore Technologies (ONT) errors are measured in relation to the ground truth of the ONT dataset, while Illumina errors are measured in relation to the ground truth for the Illumina dataset. Color scheme is capped at ±10, resulting in error greater than ±10% observing the maximum error colors. Displayed are the 20 species claiming the largest abundance in any of the Emu, minimap2, Kraken 2, Bracken, NanoCLUST, Centrifuge, or QIIME 2 results on the ONT or Illumina sample. “Other” represents the sum of all species not shown in figure for the respective column. **b** Species-level L1-norm, precision, sensitivity, and F-score for the methods evaluated in panel **a**. True positives are restricted to species with relative abundance ≥0.01% to align with guidance from ZymoBIOMICS on maximum levels of contamination.

#### Synthetic gut microbiome

Emu on ONT reads once again shows best or near-best for these metrics on the synthetic gut microbiome community. This is an intentionally challenging community containing several microbes which, even based on putative input abundance, are below 0.01% relative abundance. This is a particular form of stress-test for Emu because the EM algorithm specifically down-weights low-abundance taxa that are closely related to those in higher abundance (reflecting likelihood of sequencing error accounting for the match). Nonetheless, at the species level Emu has the best L1 distance. Additionally, Emu is only one species shy of the highest TP count but has far fewer FPs than every method aside from NanoCLUST. Although NanoCLUST does report the lowest FP counts, it also detects fewer TPs than others. Results are visualized in Supplementary Fig. 1.

### Research Application: Human Vaginal Microbiome

Variation in vaginal microbiota is associated with several urogenital diseases including bacterial vaginosis^24, 25^, a variety of sexually transmitted infections (e.g. HIV^26^), and uncategorized phenotypes such as reproductive success^24, 27^. Vaginal microbial communities can be classified into six so-called “community state types” (CSTs) I, II, III, IV-A, IV-B, and V, which are defined by relative abundance of several *Lactobacillus* species and the presence of anaerobic bacteria^28, 29^. We generated community composition from 12 vaginal samples, 6 with diagnosed bacterial vaginosis and 6 controls, using Emu and an established whole-genome shotgun (WGS) metagenomic approach. We compared CST characterizations between the two pipelines to test Emu’s ability to reproduce accepted community clusters.

#### Experimental design

12 vaginal swabs were obtained from the German Centre for Infections in Gynecology and Obstetrics at Helios Hospital Wuppertal and prepared in the Institute of Medical Microbiology, Virology, and Hospital Hygiene at the University of Duesseldorf. Samples 1-6 originate from control group patients and samples 7-12 from patients with diagnosed bacterial vaginosis. Each sample was sequenced by whole-genome and 16S ONT workflows. The whole-genome reads were processed into species- and genus-level abundance profiles using Kraken 2 and Bracken, while the 16S reads were processed with Emu.

#### 16S and WGS data comparison

Comparison of 16S and WGS sequencing data is not trivial, even when sequencing libraries are prepared from the same nucleic acid prep. Amplification and sequencing bias generally associated with marker gene sequencing approaches may occur prior to bioinformatic analysis^30^. Still, this comparison is useful to present the benefits and limitations of 16S sequencing. Since swabs contained a significant portion of host DNA (98-99% of reads classified as human by Kraken 2 and Bracken), the number of bacterial reads was lower in WGS than 16S sequencing. To reduce bias due to imbalance in sensitivity between the two methods, a species detection threshold of 0.01% was set for Emu.

Table 2 displays the most abundant bacterial genuses and four *Lactobacillus* species which are used as markers for inference of vaginal CST. 16S and WGS results are highly concordant, showing the same dominant CST-marker species across 11 of the 12 samples. Sample 10 is the only sample with differing assignment between methods. According to Bracken, it is dominated by *L. crispatus*, yet Emu shows dominance by *Megasphaera* spp. This discrepancy could be explained by the differing sequencing depths between 16S and WGS datasets. In WGS, Kraken 2 only classified as 170 reads as bacterial, whereas the 16S dataset consisted of 170,608 reads.

**Table 2.**
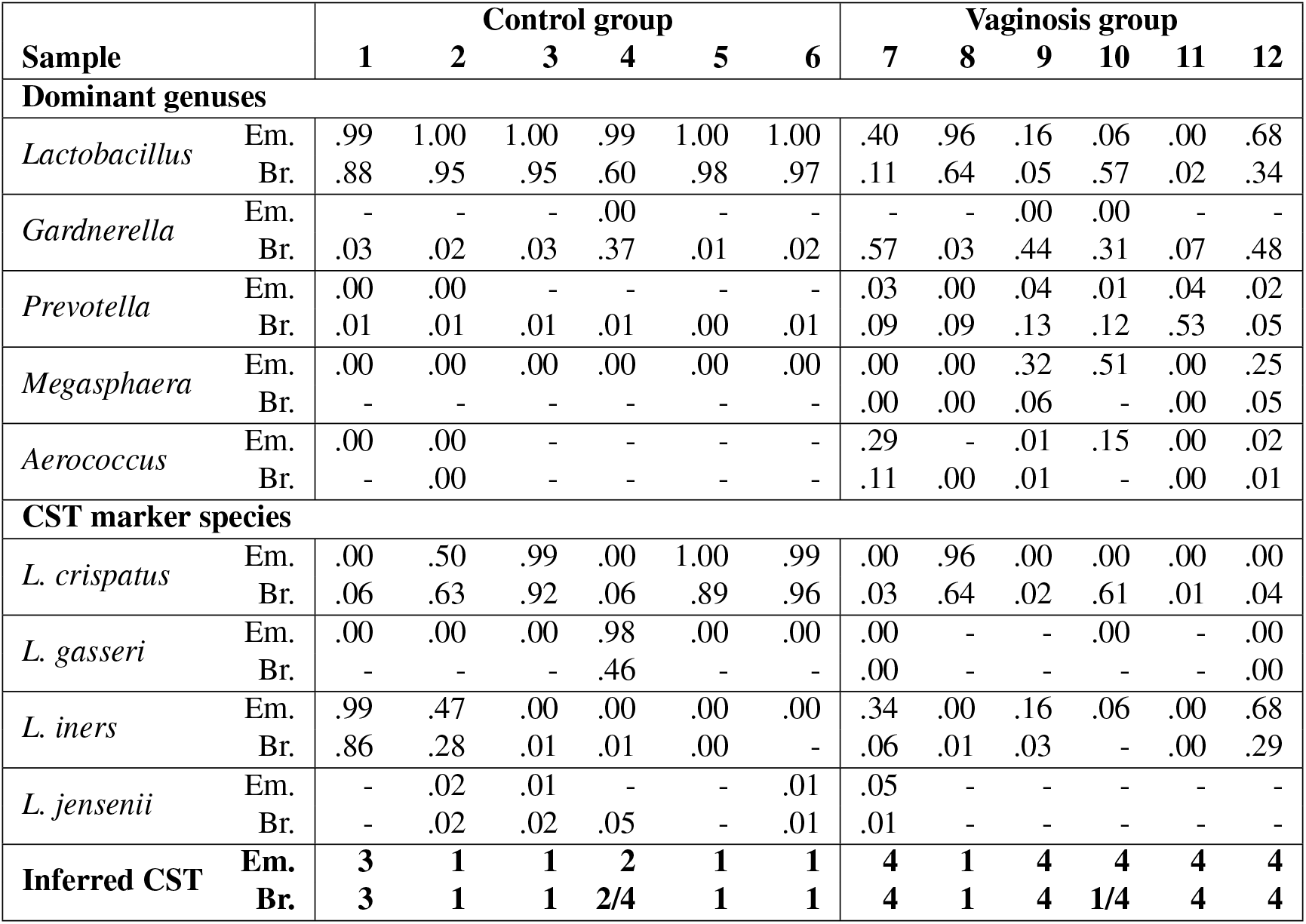
Relative abundance of dominant and marker taxons assigned by EMu from 16S rRNA ONT data and by Bracken from whole genome shotgun ONT data. Dominant genera are defined as those showing over 10% abundance in at least one sample. CST-marker species of *Lactobacillus* are defined in^28,29^. Values are rounded, so true zero values display as “-”.

| Sample |  | Control group |  |  |  |  |  | Vaginosis group |  |  |  |  |  |
| --- | --- | --- | --- | --- | --- | --- | --- | --- | --- | --- | --- | --- | --- |
|  |  | 1 | 2 | 3 | 4 | 5 | 6 | 7 | 8 | 9 | 10 | 11 | 12 |
| **Dominant genres** |  |  |  |  |  |  |  |  |  |  |  |  |  |
| *Lactobacillus* | Em. | .99 | 1.00 | 1.00 | .99 | 1.00 | 1.00 | .40 | .96 | .16 | .06 | .00 | .68 |
|  | Br. | .88 | .95 | .95 | .60 | .98 | .97 | .11 | .64 | .05 | .57 | .02 | .34 |
| *Gardnerella* | Em. | - | - | - | .00 | - | - | - | - | .00 | .00 | - | - |
|  | Br. | .03 | .02 | .03 | .37 | .01 | .02 | .57 | .03 | .44 | .31 | .07 | .48 |
| *Prevotella* | Em. | .00 | .00 | - | - | - | - | .03 | .00 | .04 | .01 | .04 | .02 |
|  | Br. | .01 | .01 | .01 | .01 | .00 | .01 | .09 | .09 | .13 | .12 | .53 | .05 |
| *Megasphaera* | Em. | .00 | .00 | .00 | .00 | .00 | .00 | .00 | .00 | .32 | .51 | .00 | .25 |
|  | Br. | - | - | - | - | - | - | .00 | .00 | .06 | - | .00 | .05 |
| *Aerococcus* | Em. | .00 | .00 | - | - | - | - | .29 | - | .01 | .15 | .00 | .02 |
|  | Br. | - | .00 | - | - | - | - | .11 | .00 | .01 | - | .00 | .01 |
| **CST marker species** |  |  |  |  |  |  |  |  |  |  |  |  |  |
| *L. crispatus* | Em. | .00 | .50 | .99 | .00 | 1.00 | .99 | .00 | .96 | .00 | .00 | .00 | .00 |
|  | Br. | .06 | .63 | .92 | .06 | .89 | .96 | .03 | .64 | .02 | .61 | .01 | .04 |
| *L. gasseri* | Em. | .00 | .00 | .00 | .98 | .00 | .00 | .00 | - | - | .00 | - | .00 |
|  | Br. | - | - | - | .46 | - | - | .00 | - | - | - | - | .00 |
| *L. iners* | Em. | .99 | .47 | .00 | .00 | .00 | .00 | .34 | .00 | .16 | .06 | .00 | .68 |
|  | Br. | .86 | .28 | .01 | .01 | .00 | - | .06 | .01 | .03 | - | .00 | .29 |
| *L. jensenii* | Em. | - | .02 | .01 | - | - | .01 | .05 | - | - | - | - | - |
|  | Br. | - | .02 | .02 | .05 | - | .01 | .01 | - | - | - | - | - |
| **Inferred CST** | **Em.** | **3** | **1** | **1** | **2** | **1** | **1** | **4** | **1** | **4** | **4** | **4** | **4** |
|  | **Br.** | **3** | **1** | **1** | **2/4** | **1** | **1** | **4** | **1** | **4** | **1/4** | **4** | **4** |

Previous literature claims healthy vaginal microbial communities to be dominated by *Lactobacillus* species^27, 28^, which aligns with the estimates conducted by Emu. Vaginal dysbiosis, on the other hand, has been associated with high abundance of generas *Gardnerella*, *Prevotella*, *Megasphaera*, *Aerococcus*^31^.

The most notable discrepancy between WGS and 16S amongst these genera is the relative abundance of *G. vaginalis*, where WGS depicts this species in significant higher abundance while 16S misses it almost altogether. This is a result of the same primer mismatch problem noted earlier for the family Bifidobacteriaceae, which *G. vaginalis* belongs to. Even with this bias, the inference community state between the Emu and Bracken workflows is consistent across samples. A heatmap displaying complete abundance profiles produced by both pipelines for each of the 12 samples is shown in the Supplementary Fig 2.

## Discussion

Of the software tools in our 16S comparisons, only NanoCLUST and Emu are designed for species-level abundance estimation using full-length 16S reads, so they should be expected to perform best in this analysis. In both mock communities, ONT reads are able to deliver notably lower L1 distances than that of Illumina reads at the species level. While this difference in L1 decreases when moving up taxonomic ranks, lower L1 distances are still achieved by ONT reads when compared to Illumina at the genus level. This highlights the effectiveness of full-length 16S reads over short read 16S analysis, as the entire 16S gene is utilized rather than restricting sequences to a subset of the hypervariable regions.

An especially notable strength of Emu is its ability to correct for initial misclassification due to ONT sequencing through the expectation-maximization algorithm. To see this we can compare Emu’s abundance estimates to minimap2. Emu returns lower L1 distance and significantly fewer false positives, at both the genus- and species-levels, across all three communities.

This error correction step is especially powerful when it comes to distinguishing between very closely related species, where the high error rate of the ONT sequences are most likely to result in confusion. The *Bacillus* species in our ZymoBIOMICS mock community illustate this situation. The species that is present as per the manufacturer is *B. subtilis*, but *B. halotolerans* differs from it by fewer than 15 bases over the length of the 16S gene. With 10-15% error in ONT reads a healthy fraction of reads map almost equally well to either one. Specifically, Of the reads claiming a minimap2 primary alignment to a *Bacillus* species, roughly 75% are correctly assigned to *B. subtilis*, 20% to *B. halotolerans*, and 5% are distributed amongst 19 other species. Fig. 3 shows how the EM algorithm addresses this as it iteratively reconsiders the statistical evidence by alternately reweighting mapping scores and abundance estimates. In the final estimate, Emu estimates 96% of the Bacillus reads as *B. subtilis*, while only falsely identifying 4 Bacillus species to account for the remaining 4%. Initially, several *Bacillus* species are inaccurately identified, but the majority are corrected after 10 iterations. It is also important to note that while minimap2 underestimates *B. subtilis* in the sample, every other method tested does so to an even greater extent.

**Figure 3.**
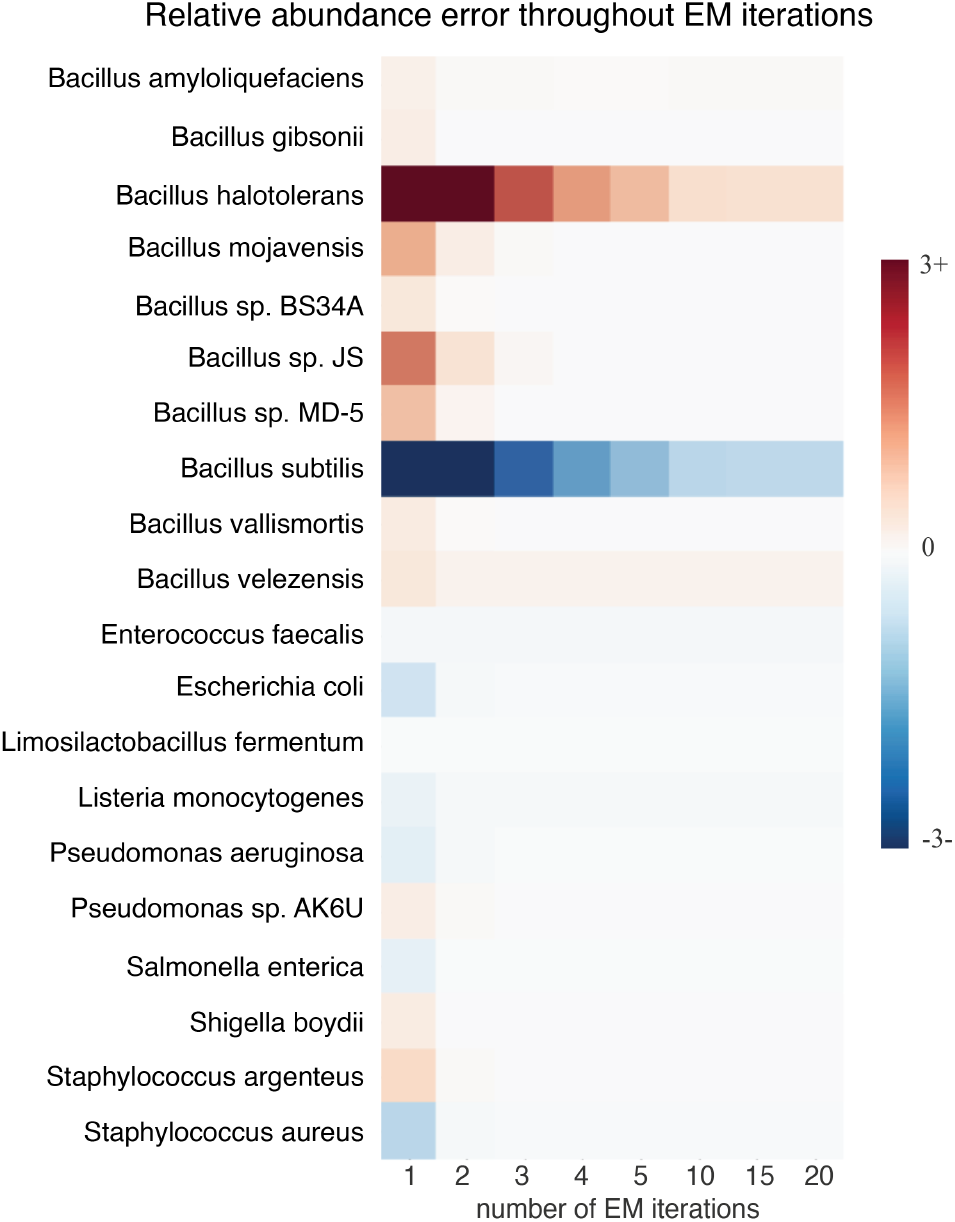
Relative error after consecutive expectation-maximization (EM) iterations on ZymoBIOMICS ONT reads. Relative error of the Emu algorithm after 1, 2, 3, 4, 5, 10, 15, and 20 EM iterations on our ZymoBIOMICS sample sequenced by a Oxford Nanopore Technologies device. The 20 most abundant species in the computational estimate are displayed. X-axis denotes the number of completed EM iterations for the results portrayed in the respective column. Darker blue represents an underestimate by the method, while darker red represents an overestimate. Color scheme is capped at ±3, resulting in error greater than ±3% observing the maximum error color.

Due to the nature of probabilistic models, the EM method generates a long tail of species with extremely low abundances. To avoid this long inaccurate list in the results, the built-in threshold for Emu is the equivalent abundance of 1 read for samples with less than 10,000 reads, and 10 reads for anything larger, meaning that for microbes truly present in a sample at lower abundance than this, Emu will not be expected to detect it. This occurs in our synthetic gut mock community where Emu does not detect *C. leptum* since our best estimate of the ground truth is that only 5 reads are present. While this abundance threshold greatly reduces false positives reported by Emu, for use cases where identification of very low abundance taxa is the priority it could be a limitation. All this to say that, of course, the desired sensitivity of the community profile is an important consideration when selecting an appropriate tool.

Additionally, since Emu is a full-length alignment-based approach, more computational resources (memory and time) are required compared to alternative methods. This is highlighted in Supplementary Table 2, and may prevent certain users from incorporating Emu into their pipeline depending on access to appropriate computing resources.

Finally, these results contain a subtle but important lesson about the practical impact false positives can have for researchers when it comes to ecological dynamics of a community. Given an environment and its resident microbiome, a common question is whether the individual microbes are strictly competing for the same resources (as plants might do for sunlight, for example) or whether there is a more complex dynamic, such as mutualism or a predator-prey relationship (as with animals). In the former case, we tend to see a small number of species or even a single highly dominant member. Yet in the latter, more diversity is generally observed and mutualistic bacteria will co-occur frequently; in the human nasal microbiome, *D. pigrim* and a species of *Cornybacterium* operate in this way^32^. In this case, a systematic tendency to overstate the number of organisms present, or to confuse a single taxon for several closely related ones, could lead to a qualitatively different understanding of the functional interactions at work.

Among the *Lactobacillus* CST-marker species in the vaginal samples, Emu suggests a single species tends to comprise over 98% of the total, whereas Bracken indicates the norm is for multiple *Lactobacillus* species to coexist. Co-occurence patterns have real significance for health researchers; one hypothesis is that a community with multiple anchoring *Lactobacilli* contains functional redundancy and thus resiliency to perturbation^29^. But if sequencing error is likely to cause one microbe to appear as a mixture of two, the resulting data could be grossly misleading. The *Bacillus* species in the ZymoBIOMICS community demonstrate the same pitfall.

The potential for long, single molecule reads to deliver higher-resolution pictures of microbial communities remains enticing, but the high rate of sequencing error has presented a formidable obstacle. Specifically, while short reads are constrained in sensitivity below the genus level, long reads are not: their difficulty is with specificity. In the case of long-reads applied to 16S amplicon sequencing, Emu represents an important improvement in minimizing this trade-off and has the potential to show the communities of well-studied environments in a new light.

In conclusion, Emu is the current top performer for species-level profiling from full-length nanopore reads and facilitates accurate characterization of microbial community composition from 16S rRNA genes.

## Methods

### Overview of Emu algorithm

To generate an accurate microbial community composition estimate from noisy full-length 16S reads, an expectation-maximization (EM) algorithm with a composition-dependent prior is developed in Emu. The algorithm starts with a frequency vector *F*, which contains the proportion of the sample that is dedicated to each species in the database. With each iteration of the algorithm, vector *F* is redistributed to increase the likelihood of the estimate. This process continues until *F* achieves only marginal gains with each proceeding cycle. At this point, vector *F* is trimmed to remove low-frequency species, then redistributed for one final time, and returned as the sample composition e stimate. Fig. 4 contains a walk through of the entire Emu algorithm, as well as a toy example.

**Figure 4.**
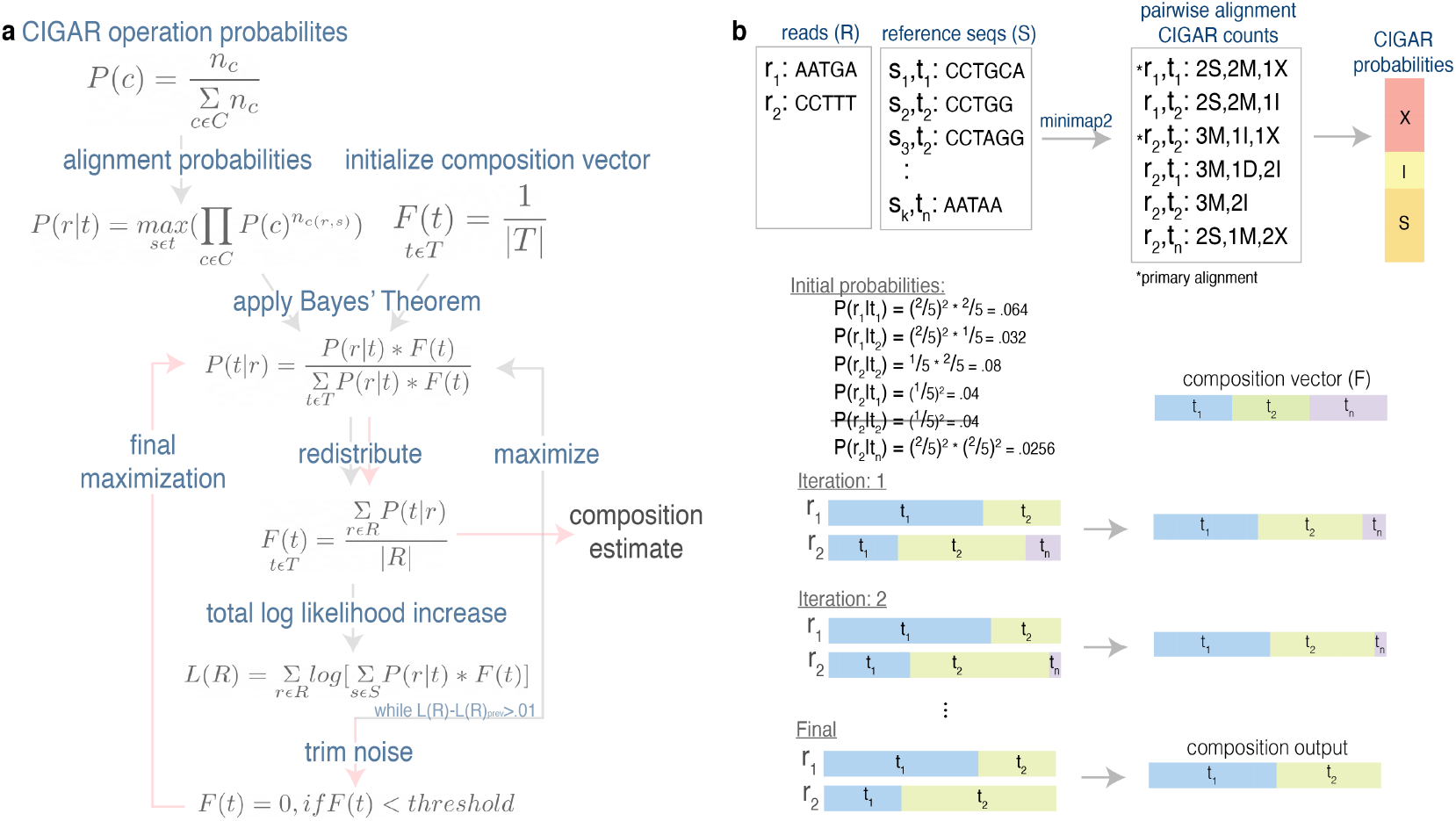
Pictorial representation of Emu algorithm. **a** The complete Emu algorithm. Follow the grey-arrowed path until expectation-maximization (EM) iterations are complete, then pink arrows are followed to the final composition estimate. The method starts by establishing probabilities for each alignment type *C =[X (mismatch), I*(*insertion*)*, S*(*so f tclipping*)], through occurrence counts in the primary alignments. Next, alignment probability *P*(*r t*) is calculated for each read, taxonomy pair (*r, t*) by assuming the maximum alignment probability between *r* and *t*. Meanwhile, an evenly-distributed composition vector *F* is initialized. The EM phase is entered by determining *P*(*t r*), the probability that *r* emanated from *t*, for all *P*(*r t*). *F* is updated accordingly and the total log likelihood of the estimate is calculated. If the total log likelihood is a significant increase over the previous iteration (>.01), then EM iterations continue. Otherwise, the loop is exited, and *F* is trimmed to remove all entries less than the set threshold. Now following the pink arrows, one final round of estimation is completed with the trimmed *F* to produce the final sample composition estimate. **b** A toy example of sample composition estimation by Emu. Sample reads *R* are mapped to database reference sequences *S* with minimap2. Utilizing the CIGAR counts in the primary alignments, nucleotide alignment probabilities 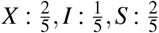 are f2ormed. The maximum alignment probability for each *r, t* is kept, and evenly-distributed vector *F* is initialized. Sample composition *F* is redistributed in each EM iteration, until the algorithm completes and returns *F* as the final relative abundance estimation.

#### Initial probabilities

To apply the EM framework, we need: 1.an initial sample composition estimate for vector *F*, and 2.alignment likelihoods *P*(*r│s*) between each sample read *r* and database reference sequence *sɛS*. Since we do not have any pre-existing knowledge regarding the sample composition, *F* starts as an evenly distributed vector 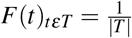, where *T* is the set of all taxonomy identifications in *S*. To identify alignment likelihoods *P*(*r│s*), we start by generating pairwise sequence alignments between *rɛR* and *sɛS* with minimap2, where *R* represents all reads in the sample. We determine the likelihood for nucleotide alignment types: mismatch (X), insertion (I), and softclippings (S), by counting the number of occurrences of each nucleotide alignment type in the primary alignments. We define these probabilities with a simple proportion: 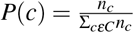, where C=[X,I,S] and *n_c_* is the sum of occurrences of that type amongst all the primary alignments. Note that excluding deletions and matches from the alignment probabilities is an empirical decision that was made after initial experimentation. This trial conducted by first ignoring all deletions to guarantee equal length between all alignments for a given read. Then, matches were removed as well, to avoid penalizing alignments for matching nucleotides.

Now that the likelihood for each type of the nucleotide alignment is defined, the likelihood for each pairwise alignment *rɛR*, *sɛS* is calculated as 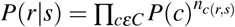, where *n_c(r,s)_* is the number of alignment type *c* observed in the alignment between *r* and *s*. In the event that no alignment is generated between *r* and *s*, then *P*(*r│s*) = 0. Since we are interested in the most-likely taxonomy of *r* rather than the most-similar sequence *s*, we keep only the highest *P*(*r│s*) for all *s* with species-level taxonomy identification *t*. Thus, the alignment probability between each read *r* and species taxonomy id *t* is calculated with 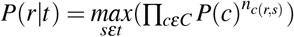 where *sɛt* represents all *s* with taxonomy id *t*. With initial probabilities set in place, the can now improve our sample composition estimation with an EM probabilistic model.

#### Redistribute sample composition

The likelihood *r* emanated from species *t* is constructed for each *P*(*r*│*t*) using Bayes’ Theorem, 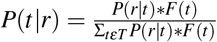. With these probabilities, *F* is redistributes as 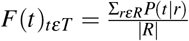. The accuracy of this estimate is evaluated by total log likelihood, *L(R) = Σ_rɛR_log[Σ_sɛS_P(r│t)*F(t)]*, which increases with each iteration. If this *L*(*R*) improvement from the previous iteration is substantial (> .01), then this re-estimation step is repeated with the updated *F*. Otherwise, redistribution is complete, and we move to the final phase of the algorithm.

#### Trim noise for final estimation

Due to the nature of the probabilistic structure in an EM model, vector *F* is likely to contain a long tail of species claiming low abundance. To avoid this long list of false positives in the output, any abundance below the set threshold will be modified to 0 at this stage. The default threshold for Emu is the abundance equivalent to 1 read for samples with under 1,000 reads and 10 reads for larger samples; however, the user can modify this parameter. Once *F* is trimmed, Emu enters one final round of abundance redistribution. The resulting final *F* is exposed as the sample composition estimation.

### Simulated read generation

Just under one million ONT reads were simulated using DeepSimulator^33^, with default settings on a synthetic metagenomic community structure. The composition was established following the composition of published mock community MBARC-26^22^. Reference 16S rRNA sequences were obtained from 16S RefSeq nucleotide sequence records^34^. For strains not present in the RefSeq 16s rRNA sequence database, all strains under the same species as the desired strain were used instead.

### Creation of gut microbiome mock community

Each gut bacterium was activated and proprogated individually in brain heart infusion (BHI) medium supplemented with hemin (5 mg/L) and yeast extract (10 g/L). Plate counting method was used to determine viable cells of cultures after 4 hours of anaerobic cultivation at 37 °C; all bacterial strains were combined with equal volume of 100 *μ*L. Cultures were then centrifuged at 12,000 g for 10 min before extra bacterial lysis with lysozyme followed by DNA extraction using MasterPure™ Complete DNA and RNA Purification Kit. DNA was quantified by Qubit kit.

### Sequencing mock communities

#### ZymoBIOMICS

Detailed description of steps taken to sequence the ZymoBIOMICS sample can be found in the **Materials and Methods** section of the study which produced these sequences^23^.

#### Synthetic gut mock

Library construction and sequencing of V4 region of the 16S ribosomal RNA gene were performed using the NEXTflex 16S V4 Amplicon-Seq Kit 2.0 (Bioo Scientific, Austin, TX) with 20 ng of input DNA, and sequences were generated on the Illumina MiSeq platform (Illumina, San Diego, CA).

Library construction and sequencing of the full-length 16S gene were performed with MinION nanopore sequencer (Oxford Nanopore Technologies, Oxford, UK) and 16S Barcoding Kit 1-24 (SQK-16S024, Oxford Nanopore Technologies, Oxford, UK).The PCR amplification and barcoding was completed with 15 ng of template DNA added to the LongAmp Hot Start Taq 2X Master Mix (New England Biolabs, Ipswich, MA). Initial denaturation at 95°C was followed by 35 cycles of 20 s at 95°C, 30 s at 55°C, 2 min at 65°C, and a final extension step of 5 min at 65°C. Purification of the barcoded amplicons was performed using the AMPure XP Beads (Beckman Coulter, Brea, CA) as per ONT’s instructions. Samples were then quantified using Qubit fluorometer (Life Technologies, Carlsbad, CA) and pooled in equimolar ratio to a total of 50-100 ng in 10 *μ*l. The pooled library was then loaded into an R9.4.1 flow cell and run as per the manufacturer’s instructions. The MINKNOW software 19.12.5 was used for data acquisition.

### Emu 16S database

The default database of Emu is a combination of rrnDB version 5.6^35^ and NCBI 16S RefSeq downloaded on September 17, 2020^34^. Duplicate species-level entries, defined as entries with identical sequences and species-level identification, were removed. The resulting database contains 49,301 sequences from 17,555 unique microbial species. Database taxonomy was also retrieved from NCBI on the same date as the RefSeq download. This database can be reproduced by utilizing the build custom database option in Emu on both the rrnDB and RefSeq sequences separately, then concatenating the results.

This combination of two databases was constructed since established databases proved insufficient for species-level classification of full-length 16S reads. This insufficiency is shown through an analysis comparing abundance estimation results calculated by Bracken with 5 different databases: the three 16S databases built into Kraken 2 (Greengenes, RDP, and SILVA), the custom Emu database described above, and the standard default database of Kraken. First, classification results for our ZymoBIOMICS ONT reads were generated with the four 16S databases through Kraken 2 as well as the standard Kraken database through Kraken. Then Bracken was utilized for abundance estimation of each of the five classification results, with defined read-length 1500. Emu default database accuracy outperformed each of the other databases tested at both the genus- and species-level in terms of L1 distance. While nearly all reads were classified at the species-level with both the Emu and Kraken default databases, RDP and SILVA were unable to make any calls at this low taxonomic rank. Full results regarding accuracy and number of reads classified are expressed in Supplementary Table 3.

### 16S quantitative comparison

Prior to each software analysis, barcodes were removed from each mock community dataset. For our two ONT datasets, the trim_barcodes function in Guppy Basecalling Software v4.4.2^36^ was applied to align with the default method of ONT sequencers. For our two Illumina datasets, Trimmomatic v0.39 was used instead. Since each method requires different formatting for database retrieval, the default database of Emu was built for each software per instructions by the developers. Reads categorized as “unclassified” were removed prior to calculating relative abundance for each method. Supplementary Note 1 contains a detailed list of all commands used.

#### Minimap2

Since minimap2 is a sequence alignment program, we converted results to classifications by restricting the minimap2 v2.17 output to only the top database hit for each input read, and classifying the read as such. Preset option for ONT was utilized for our long-read data, while the genomic short-read mapping preset was used for our Illumina data. Subsequently, taxonomy counts were generated from the primary alignment output, and relative abundance of each species was calculated as the number of species occurrences divided by the total number of aligned reads.

#### Kraken 2

Kraken 2 v2.1.1 was used to generate a custom database matching our Emu default, then ultimately produce classification results. To calculate relative abundance from Kraken 2 classification, the “clade counts” column from the Kraken 2 report (kreport) was utilized. For species-level results, only rows with “rank:S” were kept. Relative abundance for each species was then defined as the “clade counts” for that species divided by the total number of “clade counts” in the reduced kreport. This process is then repeated at the genus-level by restricting the kreport to only those rows with “rank:G”.

#### Bracken

Since Bracken was designed for abundance estimation of Kraken results, Bracken v2.5.0 was used to gather microbial abundance estimates of our Kraken 2 results described above. For full-length ONT reads, our custom Kraken 2 database was converted to a Bracken database with read lengths 1,500. The same process was applied for Illumina data, except read lengths 250 and 300 were used for ZymoBIOMICS and synthetic gut microbiome mock, respectively. Bracken abundance estimations were then generated for each dataset at the genus and species level.

#### NanoCLUST

Since NanoCLUST utilizes a BLAST database, a custom BLAST database was created to match our Emu default database. NanoCLUST v1.9 was then run on each of our long-read samples with the docker profile option. Since NanoCLUST generates relative abundance estimates at each taxonomic rank by default, no further processing was necessary.

#### Centrifuge

Centrifuge v1.0.4 was used to build a custom database and generate taxonomic classification of our 3 long-read samples. The kreport generation functionality within Centrifuge was then incorporated to create Kraken-style reports for each Centrifuge classification result. Genus- and species-level relative abundance results were calculated from these kreports in the same manor as Kraken 2 results described above.

#### QIIME 2

QIIME 2 results were produced with the classify-sklearn Naive Bayes classifier workflow of QIIME 2 2020.11.1. First, a QIIME 2 artifact representation of the default Emu database was generated with the appropriate QIIME 2 import command. Then, reference sequences were extracted appropriately based on the primer used for each sample and fit to the reference taxonomy to produce a QIIME 2 classifier. The already demultiplexed sample reads were denoised (Illumina) or dereplicated (ONT), and then classified with the appropriate pre-fit classifier. The taxonomic classifications were then collapsed to genus and species levels, and relative abundances were calculated separately for the two taxonomic rank results.

#### Establishing ground truth

Since the true microbial composition of each sequencing output is unknown, we used the following methods to establish a ground truth for our non-simulated data sets. First, our ZymoBIOMICS community. The ZymoBIOMICS Microbial Community DNA Standard used to create our ZymoBIOMICS datasets contains eight bacterial species of which assembled 16S reference genomes are provided. Thus, a “Zymo-exclusive” database containing only the assembled reference genomes for these eight species was created. The ZymoBIOMICS samples were then mapped (BWA-MEM for short-read, minimap2 for long-read) to this “Zymo-exclusive” database for accurate classification of each read. Mirroring our minimap2 workflow, reads were classified as the top hit and relative abundance was derived from these results.

A similar workflow was used to establish a ground truth for our synthetic gut microbiome community. This community was created by combining strains from 20 unique bacterial and archaeal species. However, whole-genome sequencing and de novo assembly of the isolated *Entercoccus faecium* strains, confirmed contamination with *Entercoccus faecalis*. Thus, the restricted database for this sample is limited to only the 21 species known to be in the sample, of which reference sequences were retrieved from NCBI 16S RefSeq, resulting in 45 sequences and 20 of the 21 species. However, since *Romboutsia hominis* is present in the sample, but not RefSeq, the sequence for the 16S gene of *Romboutsia hominis* was selected from GenBank^37^, and included in the exclusive database. Mapping, classification, and sample composition calculation follow the workflow for establishing ground truth of the ZymoBIOMICS community. It is important to note that this community is subject to other undocumented contamination. Supplementary Table 1 lists the established microbial compositions for each of our samples.

#### Accuracy evaluation metrics

L1-norm, or the distance between the true and inferred sample, is used to evaluate sample composition accuracy for each software. This equation can be summarized as Σ*_sɛS_│E_s_-I_s_*│, where set *S* consists of the union between all the species in the database and ground truth, and *E_s_* and *I_s_* are the respective expected and inferred relative abundances for species *s*. A perfect L1 distance is 0, while an entirely inaccurate sample composition estimate would return a L1 distance of 200. In addition, we evaluate the performance of each software with precision, sensitivity, and F-score. For this explanation, TP represents true positives, FP represents false positives, and FN represents false negatives. Precision states the proportion of claimed true positives that are truly present in the sample: 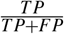. Sensitivity expresses the percentage of expected positives that were detected by the software: 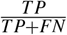. The F-score is simply the harmonic mean between the two values: 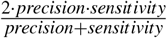. Since the ZymoBIOMICS sample is guaranteed to contain <0.01% foreign microbial DNA, all ZymoBIOMICS results are trimmed to include only taxa with abundance ≥ 0.01%, prior to calculated performance metrics.

#### Computational resources

All software analysis was completed on a Ubuntu 18.04.4 LTS system. The /usr/bin/time command was used to gather time and memory statistics. Reported CPU time is calculated by summing the user and sys time, and RAM requirements with the maximum resident size. The only except is NanoCLUST, where, computational requirements were extracted from the Nextflow execution report instead. Here, run time was gathered from the “CPU-Hours” output, and maximum resident set size from the maximum RAM reported in the “Memory” section. Computational requirements recorded for Bracken is an accumulation of both the Bracken and Kraken 2 commands, since both are required to produce the Bracken abundance estimation. Computational requirements for the QIIME 2 workflow are left out of this analysis as QIIME 2 involves several commands, making for an uneven comparison.

### Clinical vaginal samples

#### Data generation

Total DNA and RNA was extracted using ZymoBIOMICS DNA/RNA Miniprep Kit R2002. 16S Nanopore sequencing library was prepared from 10 ng of total DNA using 16S Barcoding Kit SQK-RAB204. Whole genome Nanopore library was prepared from remaining total DNA using Native Barcoding Expansion 1-12 (PCR-free) Kit EXP-NBD104 and Ligation Sequencing Kit SQK-LSK109. Data was sequenced on MinION flow cells type R9 (FLO-MIN106D) in two runs (16S run and whole genome run). Data was aquired with MINKNOW core v. 4.0.5. Basecalling and demultiplexing was done using Guppy v. 4.0.15.

#### Data analysis and databases

Computational analysis of vaginal samples was performed on a machine with CentOS Linux release 7.9.2009. Whole genome sequencing data were analyzed with Kraken v.2.1.1 and Bracken v.2.5.

Kraken 2 database was built from a custom metagenomic database, which includes all latest complete and reference genomes derived from RefSeq database in divisions bacteria, fungi, protozoa and viral of RefSeq (state 26.12.2019). The host portion of the metagenomic database is represented by 1000 genomes project reference sequence and two well-characterized human assemblies (GCA_001524155.4 and GCA_002009925.1).

Retrieved Bracken abundances at both genus- and species-level were recalculated considering only bacteria in order to align with 16S results. Therefore, total Bracken results belonging to superkingdom “Bacteria” was assumed as 100% abundance for each sample.

Emu was run on 16S sequencing data with a species detection threshold of 0.01%. Species- and genus-level abundances were retrieved from Emu output. CSTs were inferred from abundance profile considering dominance of four marker *Lactobacillus* species.

### Data availability

All sequenced samples used in this study are publicly available on Sequence Read Achieve (SRA). Both ZymoBIOMICS data sets are under BioProject ID PRJNA587452 with SRA accessions SRR10391201 for ONT and SRR10391187 for Illumina. Our gut mock community is under BioProject ID PRJNA725207. The 12 vaginal samples used for our real-world application demonstration are uploaded under BioProject ID PRJNA723982. Additionally, our simulated sequences are publicly available on OSF under project 56UF7 (https://osf.io/g5xwr/).

Study of vaginal microbiomes was IRB-approved by the ethics committee of the Medical Faculty of Heinrich Heine University.

### Code availability

Emu and all associate code are available on GitLab: https://gitlab.com/treangenlab/emu. Emu can be installed via Bioconda: https://anaconda.org/bioconda/emu.

## Acknowledgements

This work has been supported by Jürgen Manchot Foundation and Deutsche Forschungsgemeinschaft (DFG) award #428994620, by NIH grants from NIDDK P30-DK56338, NIAID R01-AI10091401, U01-AI24290 and P01-AI152999, and NINR. Q.W. was supported in part by NIH grant R21NS106640 from National Institute for Neurological Disorders and Stroke (NINDS). T.T. was supported in part by NIH grant 1P01AI152999-01 supported by National Institute of Allergy and Infectious Diseases (NIAID). M.N. was funded by a fellowship from the National Library of Medicine Training Program in Biomedical Informatics and Data Science (T15LM007093, PI: Kavraki). Computational support and infrastructure were provided by the “Centre for Information and Media Technology” (ZIM) at the University of Düsseldorf (Germany).

## Author contributions statement

A.D. and T.T. derived the Emu concept. K.C., Q.W., and M.N. developed the software. K.C., Q.W., A.T., E.R produced results for benchmarking. P.F., E.G., W.M., S.S, Q.W., T.S, and S.V. generated sequencing data for analysis. K.C., Q.W., M.N., A.T, Q.W, E.R., A.D, and T.T. contributed to writing the manuscript. All authors read and approved the manuscript.

## Competing interests

The authors declare no competing interests.

## Notes

https://gitlab.com/treangenlab/emu

### Competing Interest Statement

The authors have declared no competing interest.

