## SUPPLEMENTARY DATA for "Emu: Species-Level Microbial Community Profiling for Full-Length Nanopore 16S Reads"

#### Contents

Supplementary Figure 1: Performance on Synthetic Gut Microbiome Mock Community 2
Supplementary Figure 2: Full Abundance Profile Comparison on Vaginal Samples 3
Supplementary Table 1: Ground Truth Abundance Profiles for Three Communities 4
a) MBARC-26 Simulated 4
b) ZymoBIOMICS Community Standard 4
c) Gut Microbiome Mock Synthetic 4
Supplementary Table 2: Computational Performance Metrics 5
Supplementary Table 3: Comparison of Database Performance using Kraken2/Bracken 5
Supplementary File 2: Complete Abundance Results for Quantitative Study 6
Supplementary Note 1: Commands Used to Run Quantitative Study 7
Trim barcodes 7
Emu v1.0.1 7
Minimap2 v.2.17 7
Kraken 2 v2.1.1 7
Bracken v2.5.0 8
NanoCLUST v1.9 8
Centrifuge v1.0.4 8
QIIME 2 v2020.11 8
Establish ground truth 10

#### Supplementary Figure 1: Performance on Synthetic Gut Microbiome Mock Community

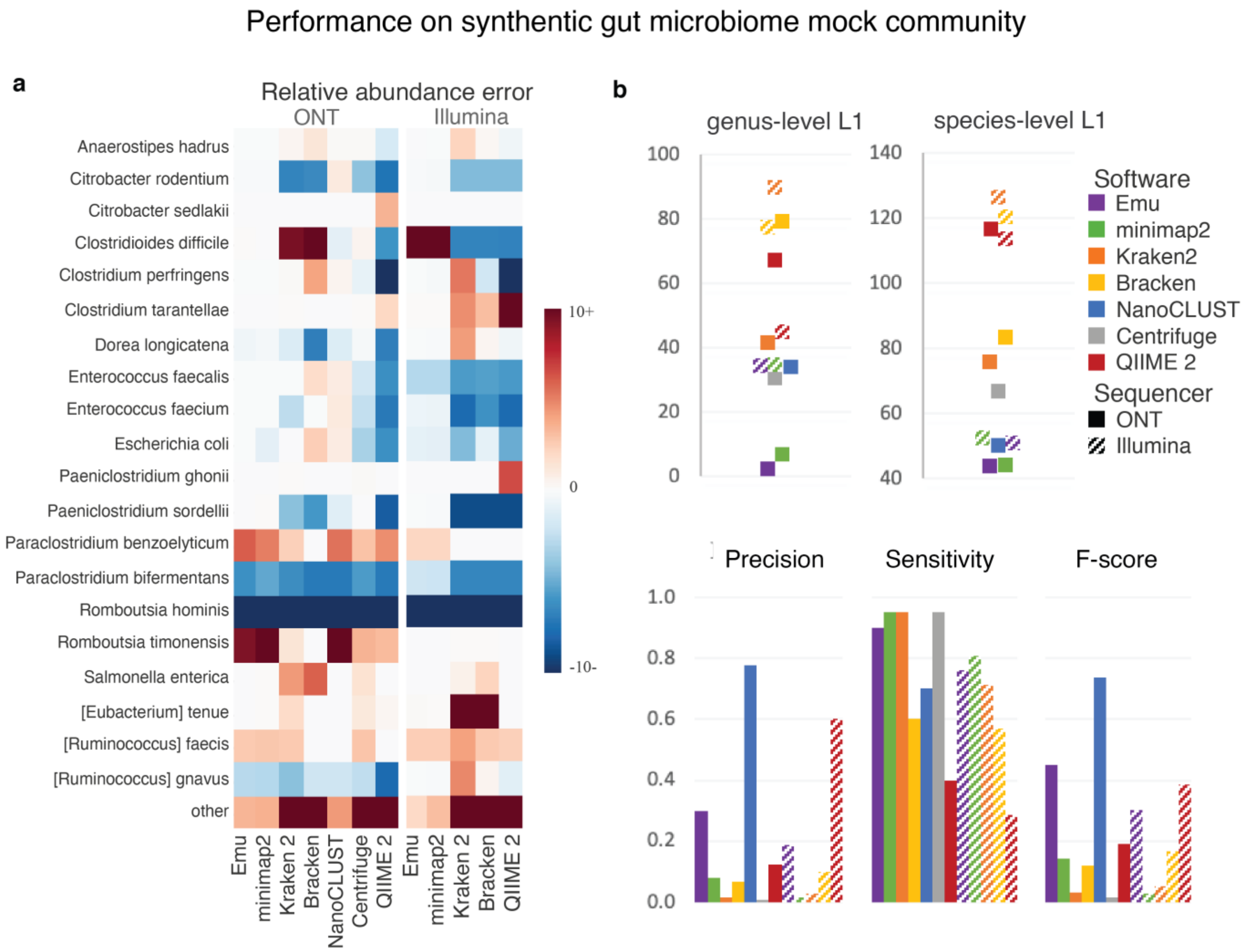


**Supplementary Figure 1. Synthetic gut microbiome mock community.** Composition accuracy for our synthetic gut microbiome mock community sequenced by Oxford Nanopore Technologies (ONT) and Illumina devices. **a** Species-level error between established ground truth abundance and estimated relative abundance. All ONT results are compared to the established truth for the ONT dataset, while all Illumina results are compared to established ground truth for the Illumina dataset. Color scheme is capped at ±10, resulting in error greater than ±10% observing the maximum error colors. Displayed are the 20 species claiming the largest abundance in any of the software results. "Other" represents the sum of all species not shown in figure for the respective column. **b** Genus-level L1-norm, as well as species-level L1-norm, precision, sensitivity, and F-score for all results expressed in panel **a**.

#### Supplementary Figure 2: Full Abundance Profile Comparison on Vaginal Samples

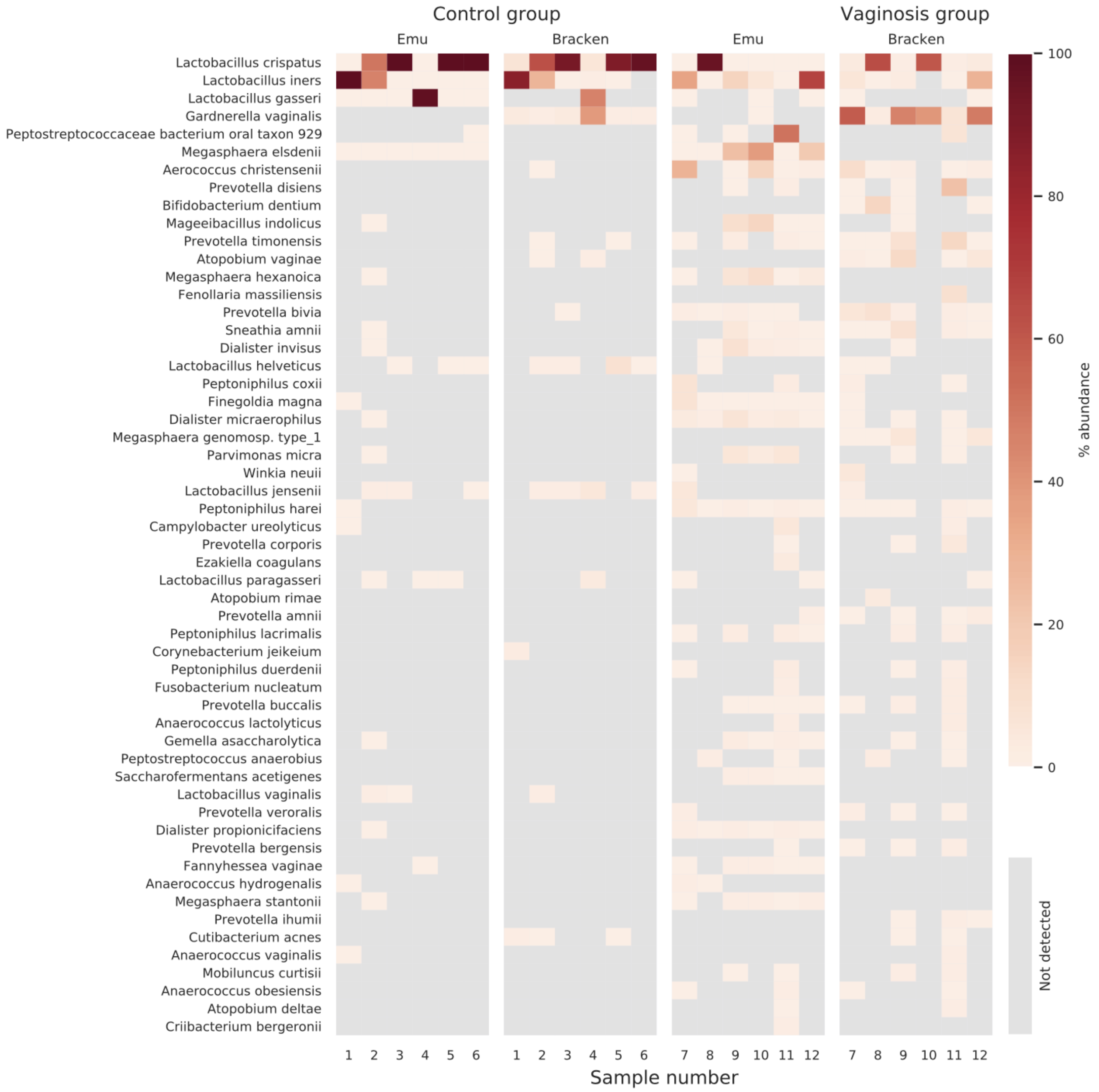


**Supplementary Figure 2. Bacterial community of 12 vaginal samples.** Species with estimated abundance of over 1% in at least one sample with either Emu or Bracken are shown. Data is grouped by condition: healthy control or vaginosis.

### Supplementary Table 1: Ground Truth Abundance Profiles for Three Communities

#### a) MBARC-26 Simulated

| Nanopore (%) | strain | NCBI Accession |
|---|---|---|
| 0.14 | Clostridium perfringens ATCC 13124 (F) | NR_121697_2 |
| 1.79 | Coraliomargarita akajimensis DSM 45221 (V) | NR_041496_1, NR_074901_1 |
| 0.13 | Corynebacterium glutamic Corynebacterium glutamicum | NR_074663_1, NR_041817_1 |
| 7.64 | Desulfotomaculum gibsoniae DSM 7213 (F) | NR_114759_1, NR_114761_1, NR_037081_1, NR_114760_1, NR_103939_1 |
| 6.99 | Desulfosporosinus acidiphilus SJ4 DSM 22704 (F) | NR_074663_1, NR_041817_1 |
| 2.58 | Desulfosporosinus meridiei DSM 13257 (F) | NR_074129_1 |
| 0.33 | Echinicola vietnamensis DSM 17526 (B) | NR_102438_1 |
| 0.10 | Escherichia coli K-12, MG1655 (P) | NR_114042_1, NR_112558_1, NR_024570_1 |
| 12.15 | Fervidobacterium pennivorans DSM 9078 (T) | NR_074097_1, NR_117583_1, NR_117584_1, NR_117582_1 |
| 7.41 | Frateuria aurantia DSM 6220 (P) | NR_074107_1, NR_040947_1 |
| 6.10 | Halovivax ruber XH-70 (E) | NR_112853_1, NR_042522_1, NR_113517_1, NR_102445_1 |
| 4.64 | Hirschia baltica ATCC 49814 (P) | NR_118878_1, NR_042123_1, NR_074121_1 |
| 0.11 | Clostridium thermocellum ATCC 27405 (F) | NR_074629_1 |
| 11.42 | Meiothermus Silvanus DSM 9946 (D) | NR_074273_1, NR_027600_1 |
| 7.85 | Natronobacterium gregoryi SP2 (E) | NR_102442_1, NR_113531_1, NR_028223_1 |
| 5.61 | Natronococcus occultus DSM 3396 (E) | NR_113534_1, NR_102453_1, NR_112871_1, NR_028255_1 |
| 0.01 | Nocardiopsis dassonvillei DSM 43111 (AT) | NR_074635_1 |
| 8.87 | Olsenella uli DSM 7084 (AT) | NR_074414_1, NR_036820_1, NR_115110_1, NR_116937_1 |
| 3.79 | Pseudomonas stutzeri RCH2 (P) | NR_113652_1, NR_116489_1, NR_114751_1, NR_118798_1, NR_103934_2 |
| 0.04 | Salmonella bongori NCTC 12419 (P) | NR_074888_1 |
| 0.17 | Salmonella enterica subsp. arizonae serovar RSK2980 ( | NR_041696_1, NR_116125_1 |
| 7.25 | Spirochaeta smaragdinae DSM 11293 (S) | NR_074741_1, NR_027585_1 |
| 3.18 | Segniliparus rotundus DSM 44985 (AT) | NR_116690_1, NR_043018_1, NR_074426_1 |
| 0.40 | Streptococcus pyogenes M1 GAS SF370 (F) | NR_112088_1, NR_028598_1 |
| 1.25 | Terriglobus roseus DSM 18391 (AD) | NR_043918_1 |
| 0.06 | Thermobacillus composti KWC4, DSM 18247 (F) | NR_041449_1, NR_102440_1 |

#### b) ZymoBIOMICS Community Standard

| theoretical (%) | Nanopore (%) | Illumina (%) | species |
|---|---|---|---|
| 17.4 | 21.7 | 19.3 | Bacillus subtilis |
| 9.9 | 12.2 | 9.0 | Enterococcus faecalis |
| 10.1 | 8.9 | 9.4 | Escherichia coli |
| 18.4 | 9.3 | 14.3 | Limosilactobacillus fermetum |
| 14.1 | 15.2 | 12.7 | Listeria monocytogenes |
| 4.2 | 1.9 | 4.8 | Psedudomonas aeruginosa |
| 10.4 | 13.0 | 13.3 | Salmonella enterica |
| 15.5 | 17.7 | 17.2 | Staphylococcus aureus |

#### c) Gut Microbiome Mock Synthetic

| Nanopore (%) | Illumina (%) | species |
|---|---|---|
| 0.060 | 0.830 | [Clostridium] innocuum |
| 0.002 | 0.001 | [Clostridium] leptum |
| 0.740 | 0.481 | [Clostridium] scindens |
| 11.011 | 7.518 | [Ruminococcus] gnavus |
| 2.108 | 3.385 | Anaerostipes hadrus |
| 0.009 | 0.756 | Bacteroides thetaiotaomicro |
| 0 | 2.047 | Bifidobacterium dentium |
| 7.286 | 4.441 | Citrobacter rodentium |
| 5.949 | 6.784 | Clostridioides difficile |
| 17.890 | 15.415 | Clostridium perfringens |
| 6.910 | 6.722 | Dorea longicatena |
| 6.192 | 5.706 | Enterococcus faecalis |
| 7.123 | 7.832 | Enterococcus faecium |
| 6.120 | 4.998 | Escherichia coli |
| 0.004 | 0.001 | Gemmiger formicilis |
| 0.594 | 0.766 | Intestinibacter bartlettii |
| 8.301 | 8.939 | Paeniclostridium sordellii |
| 7.045 | 6.596 | Paraclostridium bifermentan |
| 0.019 | 0.942 | Phocaeicola vulgatus |
| 12.176 | 12.701 | Romboutsia hominis |
| 0.462 | 3.139 | Roseburia faecis |

#### Supplementary Table 2: Computational Performance Metrics

| | | Nanopore | | | | | | Illumina | | | |
|---|---|---|---|---|---|---|---|---|---|---|---|
| | | Emu | minimap2 | Kraken 2 | Bracken | NanoCLUST | Centrifuge | Emu | minimap2 | Kraken 2 | Bracken |
| MBARC-26 | time (m) | 249 | 234 | 2 | 2 | 249 | 20 | | | | |
| | RAM (GB)* | 319 | 318 | 0 | 0 | 16 | 0 | | | | |
| Zymo | time (m) | 1048 | 886 | 3 | 3 | 204 | 38 | 62 | 53 | 0 | 0 |
| | RAM (GB) | 420 | 408 | 0 | 0 | 34 | 0 | 2 | 1 | 0 | 0 |
| Gut Mock | time (m) | 221 | 200 | 1 | 1 | 270 | 12 | 10 | 6 | 0 | 0 |
| | RAM (GB) | 373 | 368 | 0 | 0 | 34 | 0 | 1 | 1 | 0 | 1 |

*RAM usage represented maximum resident set size

#### Supplementary Table 3: Comparison of Database Performance using Kraken2/Bracken

The following table shows an evaluation of performance Kraken 2/Bracken on the ZymoBIOMICS ONT data with different databases. The middle three are those made available with Kraken 2. The rightmost column includes results of Kraken/Bracken (that is, Kraken 1) on the default database. The leftmost column is the performance on the combined rrnDB/NCBI database that was built for Emu.

**Data set**: ZymoBIOMICS (ONT) **read count**: 1,096,959

| | database<br>classifier | **Emu**<br>Kraken 2 | **Greengenes**<br>Kraken 2 | **Silva**<br>Kraken 2 | **RDP**<br>Kraken 2 | **Kraken**<br>Kraken |
|---|---|---|---|---|---|---|
| genus | TP [8] | 8 | 5 | 7 | 8 | 7 |
| | FP | 60 | 35 | 24 | 72 | 60 |
| | L1-norm | 23 | 102 | 58 | 63 | 40 |
| | precision | 0.12 | 0.13 | 0.25 | 0.09 | 0.10 |
| | sensitivity | 1.00 | 0.63 | 1.00 | 0.88 | 0.88 |
| | F-score | 0.21 | 0.21 | 0.40 | 0.16 | 0.19 |
| | reads kept* | 1,040,877 | 747,277 | 1,092,151 | 1,011,567 | 1,077,148 |
| | % reads kept | 94.9 | 68.1 | 99.6 | 92.2 | 98.2 |
| species | TP [8] | 8 | 2 | - | - | 7 |
| | FP | 191 | 44 | - | - | 171 |
| | L1-norm | 66 | 179 | - | - | 78 |
| | precision | 0.04 | 0.04 | - | - | 0.04 |
| | sensitivity | 1.00 | 0.25 | - | - | 0.88 |
| | F-score | 0.08 | 0.07 | - | - | 0.08 |
| | reads kept | 1,035,110 | 557,256 | 0 | 0 | 1,075,754 |
| | % reads kept | 94.4 | 50.8 | 0.0 | 0.0 | 98.1 |

*"new_est_reads" column sum in Bracken output

#### Supplementary File 2: Complete Abundance Results for Quantitative Study

**File:** `supp_file2_complete_abundance_results.xlsx` (included with submission)

This Excel file contains the complete abundance profile outputs for each of the methods and data sets contained in Table 1. The file has 5 sheets, each one with a self-descriptive name and corresponding to a segment of Table 1:

A) MBARC-26
B) Zymo - Nanopore
C) Zymo - Illumina
D) Gut - Nanopore
E) Gut - Illumina

In each table, the first column is labeled with the ground truth abundances. In tab (A), it is labeled “truth” since the data is simulated and ground truths are known. In tabs (B) – (E) it is labeled “theoretical” since an algorithm was used to compute a putative ground truth value for these communities.

### Supplementary Note 1: Commands Used to Run Quantitative Study

Below are commands used to generate comparison results on our 3 Oxford Nanopore Technologies (ONT) datasets and 2 Illumina datasets. The following variables are used:

```
<sample path>: path to sample(s), ONT or Illumina
<sample>: ONT reads [.fastq]
<sample f>: Illumina PE forward reads [.fastq]
<sample r>: Illumina PE reverse reads [.fastq]
<db fasta>: custom database sequences [.fasta]
<db nodes>: custom database nodes [.dmp]
<db names>: custom database names [.dmp]
<db seq2tax map>: custom database sequence id to tax id map [.txt]
```

#### Trim barcodes

ONT

*Guppy Basecalling Software v4.4.2:*

```
$ guppy_barcoder --input_path <sample_path> --save_path <output_path>
--barcode_kits <kits> --trim_barcodes
```

- Our ZymoBIOMICS sample used kits SQK-RAB201, while our gut mock sample used SQK-16S024.

Illumina

*Trimmomatic v0.39*

```
$ java -jar trimmomatic-0.39.jar PE <sample_f> <sample_r> <out_sample_f>
   <out_sample_r> ILLUMINACLIP:adapters/TruSeq3-PE.fa:2:30:10:2:keepBothReads
```

#### Emu v1.0.1

ONT

```
$ emu abundance <sample>
```

Illumina

```
$ emu abundance --type sr <sample_f> <sample_r>
```

#### Minimap2 v.2.17

The following commands were used to collect primary alignments. Further processing was completed to calculate relative abundance from results.

ONT

```
$ minimap2 -ax map-ont -N 1 <db_fasta> <sample>
```

Illumina

```
$ minimap2 -ax sr -N 1 <db_fasta> <sample_f> <sample_r>
```

#### Kraken 2 v2.1.1

Build database

```
$ kraken2-build --download-taxonomy --db <db_name>
#replace name.dmp & nodes.dmp files
$ kraken2-build --add-to-library <db_fasta> --db <db_name>
$ kraken2-build --build --db <db_name>
```

ONT

```
$ kraken2 --db <db_name> <sample> --output <out_classification> --report
   <out_kreport>
```

Illumina

```
$ kraken2 --db <db_name> <sample-f> <sample_r> --output <out_classification>
   --report <out_kreport>
```

#### Bracken v2.5.0

Build databases

```
$ bracken-build -d <KRAKEN2_DB_PATH>/<db_name> -l <read_len> -k 35 -x
   <kraken_installation>
```

- read len 1500 was used for our ONT reads, while 300 and 250 were used for our Illumina reads.

ONT

```
$ bracken -d <KRAKEN2_DB_PATH>/<db_name> -i <kraken2_kreport> -r 1500 -l [G|S] -o
   <out_file>
```

Illumina

```
$ bracken -d <KRAKEN2_DB_PATH>/<db_name> -i <kraken2_kreport> -r [250|300] -l [G|S]
   -o <out_file>
```

- read len 250 was used for our ZymoBIOMICS community, and 300 for our gut mock community.

#### NanoCLUST v1.9

Build databases

```
$ makeblastdb -in <db_fasta> -parse_seqids -blastdb_version 5 -taxid_map
   <db_seq2tax_map> -title <db_name> -dbtype prot
```

ONT

```
$ nextflow run main.nf -profile docker --reads <sample> --db <db_fasta> --tax
   <db_taxonomy_path> --outdir <out_dir>
```

#### Centrifuge v1.0.4

Build databases

```
$ centrifuge-build --conversion-table <db_seq2tax_map> --taxonomy-tree <db_nodes> -
   -name-table <db_names> <db_fasta> <db_name>
```

ONT

```
$ centrifuge -q -x <db_name> <sample> -S <out_classification> --report-file
   <out_report>
$ centrifuge-kreport -x <db_name> <out_report> > <out_kreport>
```

#### QIIME 2 v2020.11

Import database

```
$ qiime tools import --input-path <db_fasta> --type FeatureData[Sequence] --output-
   path <reference_sequences>
```

```
$ qiime tools import --input-path <db_taxonomy_path> --type FeatureData[Taxonomy] -
   -output-path <reference_taxonomy>
```

Extract reference sequences with primers

```
$ qiime feature-classifier extract-reads --i-sequences <reference_sequences> --p-f-
   primer <forward_primer> --p-r-primer <reverse_primer> --o-reads
   <reference_extracted_reads>
```

- All ONT reads used full-length 16S primers: AGAGTTTGATCMTGGCTCAG, CGGTTACCTTGTTACGACTT.
- ZymoBIOMICS Illumina reads were targeted for hypervariable regions V4-6 using primers: GTGCCAGCMGCCGCGGTAA, ACAACACGAGCTGACGAC.
- The synthetic gut mock community used the v4 region with primers: GTGCCAGCMGCCGCGGTAA, GGACTACHVGGGTWTCTAAT.

Fit classier

```
$ qiime feature-classifier fit-classifier-naive-bayes --i-reference-reads
   <reference_extracted_reads> --i-reference-taxonomy <reference_taxonomy> --o-
   classifier <classifier>
```

ONT

```
$ qiime tools import --type SampleData[SequencesWithQuality] --input-path
   <path_to_sample> --input-format CasavaOneEightSingleLanePerSampleDirFmt --
   output-path <demultiplexed_sample>
$ qiime vsearch dereplicate-sequences --i-demultiplexed-seqs <demultiplexed-sample>
   --o-dereplicated-table <sample_derep_table> --o-dereplicated-sequences
   <sample_derep_seqs>
$ qiime feature-classifier classify-sklearn --i-classifier <classifier> --i-reads
   <sample_derep_seqs> --p-n-jobs <num_threads> --o-classification
   <sample_sklearn_classification>
$ qiime taxa collapse --i-table <sample_derep_table> --i-taxonomy
   <sample_sklearn_classification> --p-level [6|7] --o-collapsed-table
   <sample_collapsed_table>
$ qiime feature-table relative-frequency --i-table <sample_collapsed_table> --o-
   relative-frequency-table <sample_collapsed_rel_frequency>
```

Illumina

```
$ qiime tools import --type SampleData[PairedEndSequencesWithQuality] --input-path
   <path_to_sample> --input-format CasavaOneEightSingleLanePerSampleDirFmt --
   output-path <demultiplexed_sample>
$ qiime dada2 denoise-paired --i-demultiplexed-seqs <demultiplexed_sample> --p-n-
   threads <num_threads> --p-trunc-len-f 0 --p-trunc-len-r 0 --o-table
   <sample_denoised_table> --o-representative-sequences <sample_denoised_sequences>
   --o-denoising-stats <sample_denoising_stats>
$ qiime feature-classifier classify-sklearn --i-classifier <classifier> --i-reads
   <sample_denoised_seqs> --p-n-jobs <num_threads> --o-classification
   <sample_sklearn_classification>
$ qiime taxa collapse --i-table <sample_denoised_table> --i-taxonomy
   <sample_sklearn_classification> --p-level [6|7] --o-collapsed-table
   <sample_collapsed_table>
$ qiime feature-table relative-frequency --i-table <sample_collapsed_table> --o-
   relative-frequency-table <sample_collapsed_rel_frequency>
```

- Taxa level 6 is used for collapsing to genus level.
- Taxa level 7 is used for collapsing to species level.

#### Establish ground truth

The following commands were used to collect primary alignments. Further processing was completed to calculate relative abundance from results. For our ZymoBIOMICS community, restricted database sequences contains all 8 assembled sequences from ZymoBIOMICS. For our gut mock community, all NCBI RefSeq sequences from the 21 expected species are included. Detailed descriptions of both processes are provided in the manuscript.

ONT

```
$ minimap2 -ax map-ont <restricted_db_sequences> <sample>
```

Illumina

```
$ bwa index <restricted_db_sequences>
$ bwa mem restricted_db_sequences> <sample_f> <sample_r>
```
